# Neuronal activity induces symmetry breaking in neurodegenerative disease spreading

**DOI:** 10.1101/2023.10.02.560495

**Authors:** Christoffer G. Alexandersen, Alain Goriely, Christian Bick

## Abstract

Dynamical systems on networks typically involve several dynamical processes evolving at different timescales. For instance, in Alzheimer’s disease, the spread of toxic protein throughout the brain not only disrupts neuronal activity but is also influenced by neuronal activity itself, establishing a feed-back loop between the fast neuronal activity and the slow protein spreading. Motivated by the case of Alzheimer’s disease, we study the multiple-timescale dynamics of a heterodimer spreading process on an adaptive network of Kuramoto oscillators. Using a minimal two-node model, we establish that heterogeneous oscillatory activity facilitates toxic outbreaks and induces symmetry breaking in the spreading patterns. We then extend the model formulation to larger networks and perform numerical simulations of the slow-fast dynamics on common network motifs and on the brain connectome. The simulations corroborate the findings from the minimal model, underscoring the significance of multiple-timescale dynamics in the modeling of neurodegenerative diseases.

## 1 Introduction

Mathematical models of dynamical processes on networks are crucial to our understanding of pandemics, economics, opinion formation, evolution, ecology, and neurodegenerative disease [1–5]. While the underlying network structure is traditionally assumed to be static, it has become clear that network adaptivity is a crucial constituent to many real-world dynamical systems [6, 7]. For example, infectious diseases spreading [8] and biological neural circuits [9] alter the network structures they are evolving on, establishing a feedback loop between dynamics and network structure. Networks exhibiting such mutual interactions between dynamical processes and network topology are referred to as adaptive, or coevolutionary, networks [6, 10, 11]. In a range of applications, including oscillator networks [12–15], consensus dynamics [16], and epidemic-resource dynamics [17], coevolutionary dynamics may also operate on disparate timescales. Although techniques from geometric singular perturbation theory [18] and averaging theory [19] can provide insights into the emerging multiple-timescale dynamics, these methods become daunting in high dimensions.

A crucial—yet poorly understood—example of an adaptive network with multiple timescale dynamics is the human brain during neurodegenerative diseases, such as Alzheimer’s disease. The defining feature of Alzheimer’s disease is the accumulation of toxic variants of amyloid-*β* and tau protein aggregates throughout the brain [20]. It is believed that these toxic variants are produced by a prion-like mechanism, where toxic variants of the protein transform healthy variants into toxic ones [21]. Although both amyloid-*β* and tau are fundamental to the disease, the presence of tau correlates more significantly with cognitive decline. Furthermore, it has been shown that tau proteins spread throughout the brain following axonal pathways [22, 23], leading to neurodegeneration and decreases in neuronal activity levels [21]. Tau proteins tend to follow a general spreading sequence—called the Braak staging pattern—starting in the entorhinal cortex. However, the basis for the initiation of Braak staging in the entorhinal cortex and the ensuing spreading pattern remains disputed. Furthermore, subgroupings of patients according to systematic aberrations in Braak staging patterns further complicate our picture of the disease [24, 25]. In recent years, however, it has become clear that neuronal activity plays a crucial role in the spreading of tau protein. Specifically, it has been shown that neurons with higher firing rates transport tau proteins at a higher rate into their neighbors [26–28]. As such, neuronal activity increases the outward transport of tau proteins, while tau proteins lower neuronal activity levels. Importantly, protein spreading and neuronal activity evolve on vastly different timescales; protein spreading operates on a timescale of years while neuronal activity operates on a timescale of seconds. The newly discovered bidirectional relationship between neuronal activity and protein spreading may be the missing link in our understanding of Alzheimer’s disease and other neurodegenerative diseases.

Mathematical modeling of neurodegenerative diseases has mostly focused on protein spreading and neuronal activity in isolation. On the one hand, the slow evolution of the protein spreading and subsequent damage to the neural networks in Alzheimer’s disease can be captured with a continuum approach using Fischer-KPP equations [29]. However, protein spreading dynamics can also be effectively modeled from a network perspective using structural network reconstructions of the human brain. This idea was initially introduced in [30]—and later expanded in [31]—and involves simplifying continuum models, easing the investigation of staging patterns [32, 33] and parameter estimation through Bayesian techniques [34]. Furthermore, a separate and successful approach employs a heterodimer model to investigate how amyloid-*β* and tau spread during Alzheimer’s disease [35, 36]. On the other hand, the activity of individual neurons that give rise to neural oscillations—which are fast relative to disease progression—are captured by models of neuronal dynamics, such as the highly detailed Hodgkin–Huxley model which can emulate pathologies by incorporating defects in ion channel conductivity. However, the Hodgkin–Huxley model becomes intractable in larger networks of neurons, where oscillator models such as Kuramoto oscillators [37], theta neurons [38], and integrate-and-fire neurons [39] have shown great utility. In these models, the instantaneous oscillator frequencies are commonly interpreted as neuronal firing rates, which is a common metric for neuronal activity. Only recently have the two aspects of slow disease progression and fast neural dynamics been captured in a single modeling framework; see for example [40, 41].

Motivated by the progression of Alzheimer’s disease, we here develop a multiple timescale approach to elucidate the dynamics of spreading processes and oscillator dynamics on adaptive networks. More specifically, we formulate a multiple timescale system where a slow heterodimer spreading process occurs on a network of fast Kuramoto oscillators. The presence of protein slows the natural frequencies of the Kuramoto oscillators, while the instantaneous frequencies of the Kuramoto oscillators increase the outward transport of protein from their respective nodes. The network structure is adaptive, as the Kuramoto frequencies alter the transport rates by scaling the link weights of the spreading network. In other words, the Kuramoto oscillators are enforcing a global adaptivity rule on the spreading process. With the goal of elucidating the role of fast oscillatory processes on the spreading patterns and vice versa, we begin by studying a minimal two-node model using slow manifold reduction and ad hoc averaging before corroborating our findings with numerical simulations of the generalized network model. Conclusively, we find that heterogeneously distributed frequencies of oscillators destabilize the spreading process by lowering the threshold for toxic outbreaks and inducing symmetry breaking in the spreading patterns.

This article is organized as follows: In Section 2, we consider the heterodimer model on a minimal network of two nodes with asymmetric link weights to reflect the effect of activity on the spreading dynamics. In Section 3, we consider a multiple timescale two-node system, now equipped with both heterodimer and Kuramoto dynamics, which we call the *heterodimer-oscillator*. In Section 4, we support the findings from the minimal heterodimer-oscillator system by performing numerical simulations on common motifs found in complex networks and investigating the effect oscillatory activity can have on tau spreading in the human brain during Alzheimer’s disease.

## 2 Heterodimer dynamics

In this section, we build on the classical heterodimer model for a simple 2-node graph and introduce asymmetry in the coupling between the nodes to understand its impact on the system dynamics. Specifically, we identify a pair of fixed points exchanging stability at a transcritical bifurcation and observe that the asymmetrical coupling not only shifts the location of this bifurcation in parameter space but also disrupts symmetries within the fixed points. The dynamical behavior of the asymmetrically-coupled heterodimer model will be instrumental in our analysis of the full system with coevolutionary spreading and oscillator dynamics later on in Section 3.

### 2.1 The heterodimer model

The heterodimer model describes a process of healthy proteins being converted into toxic proteins by a second-order rate equation. The heterodimer model is often used in the context of networks, over which both the healthy and toxic proteins are spreading. We assume that the process takes place on a network with *N* nodes defined by a weighted adjacency matrix **W** = (*W*_*ij*_). For **W** we define the standard graph Laplacian **L** = (*L*_*ij*_) with components

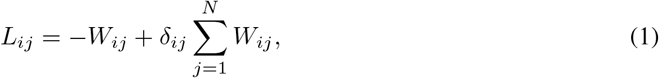

where *δ*_*ij*_ is the Kronecker symbol. According to the heterodimer model, the evolution of the concentration of healthy proteins *u*_*i*_ ≥ 0 and of toxic proteins *v*_*i*_ ≥ 0 at node *i* is given by

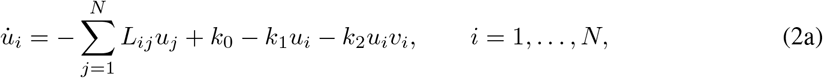

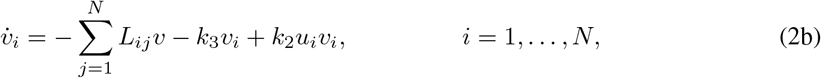

where *k*_0_ > 0 is the healthy protein production rate, *k*_1_ > 0 and *k*_3_ > 0 are the healthy and toxic clearance (protein degradation) rates, and *k*_2_ *>* 0 is the rate of conversion from healthy to toxic proteins.

With the ultimate goal of understanding how the possible dynamics of this system are affected by oscillatory activity, we start with the simple case of two nodes connected by an undirected link as shown in Figure 1(a):

**Figure 1:**
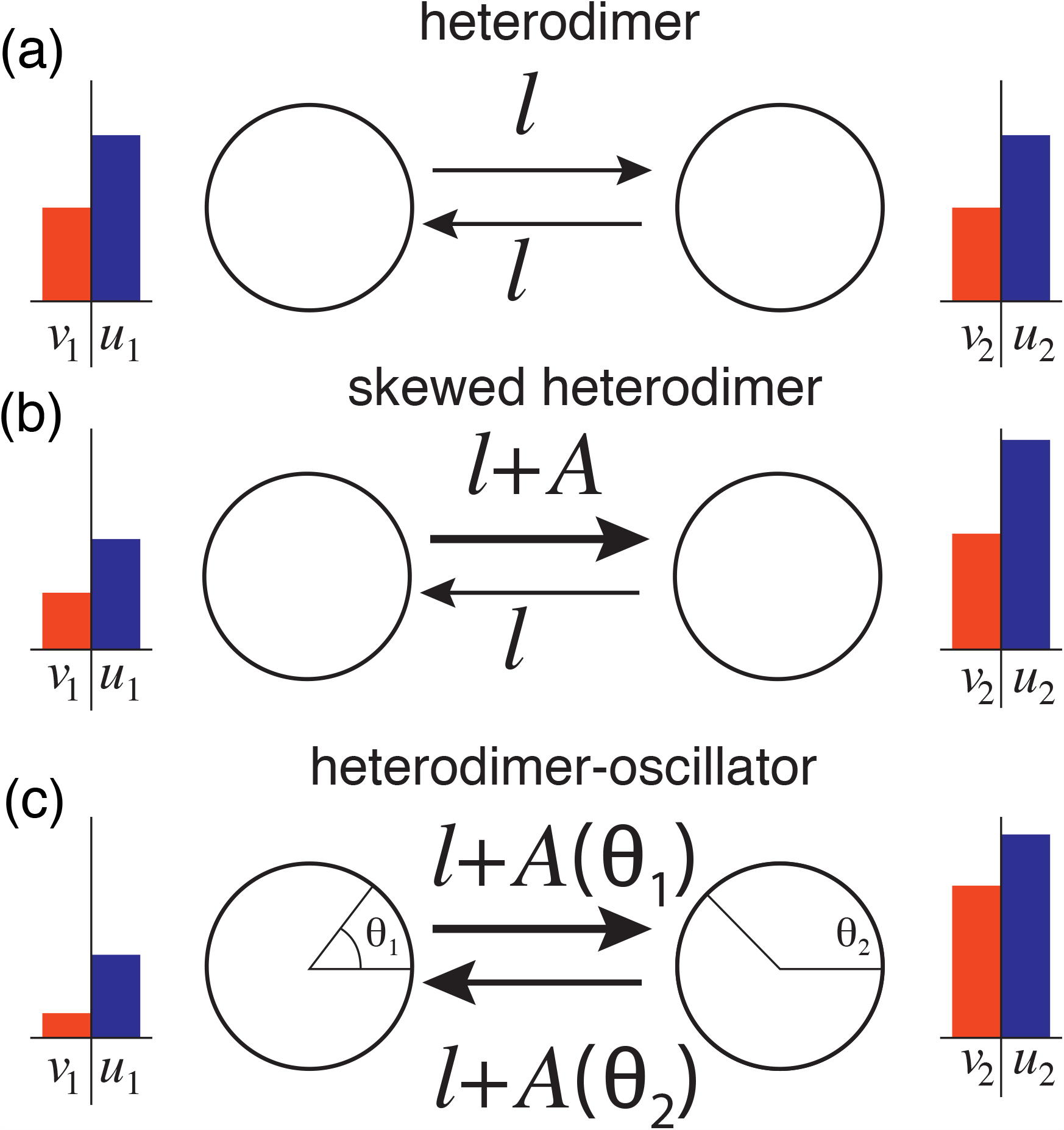
Overview of the heterodimer variations. *(a)* The original heterodimer model, with healthy and toxic species transported between the nodes at equal rates. *(b)* The skewed heterodimer model where node 1 has higher activity and thus increases the transport rate into node 2. Note that toxic species do not affect the activity parameter *A. (c)* The heterodimer-oscillator model, where each node harbors an oscillator operating at a faster time rate than the spreading process. The oscillators are coupled and their frequency determines the transport rate of species between the nodes; in this illustration, node 1 has a higher frequency. Conversely, the toxic species affect the intrinsic frequency of the oscillators.

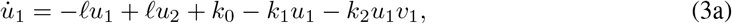

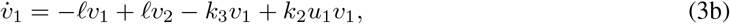

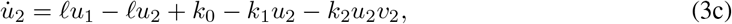

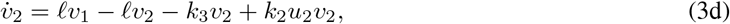

where *ℓ >* 0 is the single, reciprocal weight link. Note that all parameters and variables are nonnegative. The system has two fixed points. In general, we refer to a fixed point as *healthy* if *v*_*i*_ = 0 for all *i* = 1, …, *N* and *toxic* if *v*_*i*_ *>* 0 for at least one *i* ∈{1, …, *N*}. The 2-node heterodimer model has exactly one healthy fixed point (denoted by a superscript H) and one toxic fixed point (superscript T), given by

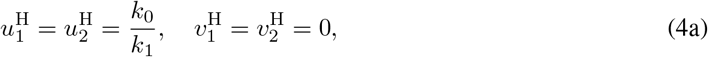

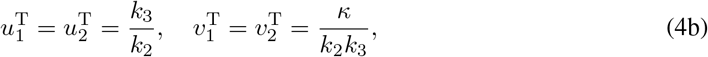

where *κ* = *k*_0_*k*_2_− *k*_1_*k*_3_.

In terms of the dynamics, we are mostly interested in the transition between healthy states and toxic states. In other words, we are interested in bifurcations where a healthy equilibrium loses stability and a toxic equilibrium becomes stable. For (3), a direct computation of the linearized system around the healthy equilibrium indicates that healthy and toxic equilibria interchange stability through a transcritical bifurcation occurring at *κ* = 0. Indeed, the stability of the healthy state is governed by a single eigenvalue

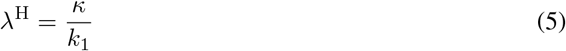

of the system’s Jacobian matrix evaluated at the healthy fixed point. Hence, we conclude that the healthy state is stable for *κ* ≤ 0 and the toxic fixed point is stable for *κ* ≥ 0.

### 2.2 The skewed heterodimer model

To understand the effect of activity dynamics on spreading, we now consider a constant activity *A*≥ 0 that affects the spreading as shown in Figure 1(b) but exclude the effect that spreading may have on activity dynamics. Assuming that the activity process *A >* 0 taking place in node 1 increases spreading to its neighbor, we obtain a *skewed heterodimer model* where the concentrations evolve according to

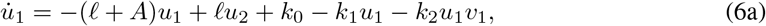

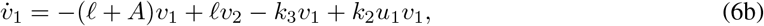

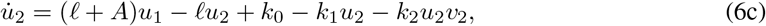

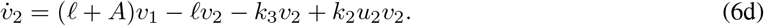

If *A* = 0 we recover (3). For the skewed heterodimer model (6), there is a single healthy fixed point

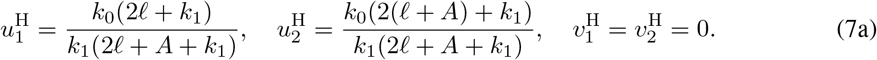

Note that introducing *A* breaks the symmetry in the healthy fixed point between the two nodes, which previously were independent of *ℓ*. Eliminating *u*_1_, *u*_2_ and *v*_1_ from the first fixed points, we find a cubic equation for the toxic fixed point 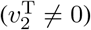 given by

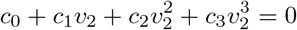

with coefficient values given in Appendix A.

To identify transitions between healthy and toxic states, we linearize the vector field at the healthy fixed point. The eigenvalues of the Jacobian at the healthy fixed point are

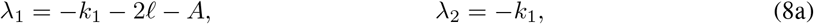

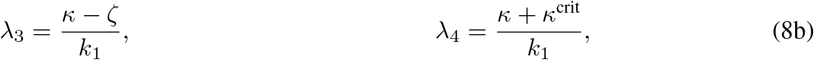

where *κ*^crit^ and *ζ* are given by

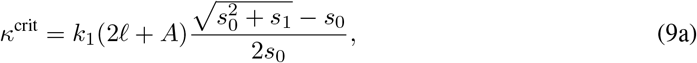

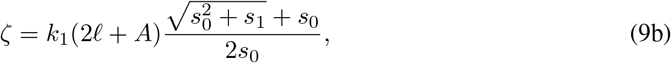

with constants *s*_0_ = *k*_1_(2*ℓ* + *A*)(2*ℓ* + *A* + *k*_1_), *s*_1_ = 4*A*^2^*k*_0_*k*_2_(*k*_1_(2*ℓ* + *A* + *k*_1_) + *k*_0_*k*_2_). Since all parameters are positive, it follows that *κ*^crit^ ≥0 and *ζ* ≥ 0. As such, we have that *λ*_3_ ≤ *λ*_4_ and *λ*_1_ *< λ*_2_ *<* 0, and hence *λ*_4_ dictates the stability of the healthy fixed point. The fixed point switches stability at a critical value *κ* =−*κ*^crit^. Freezing all parameters but the toxic clearance rate *k*_3_, we look at the bifurcation in terms of the parameter *k*_3_, with critical value

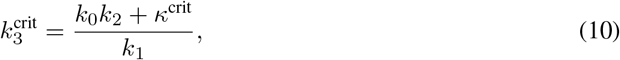

which satisfies 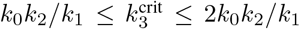 and is monotonically increasing in *A* (see Appendix B). We conclude that introducing the activity parameter *A* shifts the transcritical bifurcation to higher values with respect to *k*_3_. The effect of activity is to *destabilize* the healthy fixed point as shown in Figure 2. Equivalently, in terms of neuroscientific applications, heterogeneous neuronal activity pushes neurons toward pathology.

**Figure 2:**
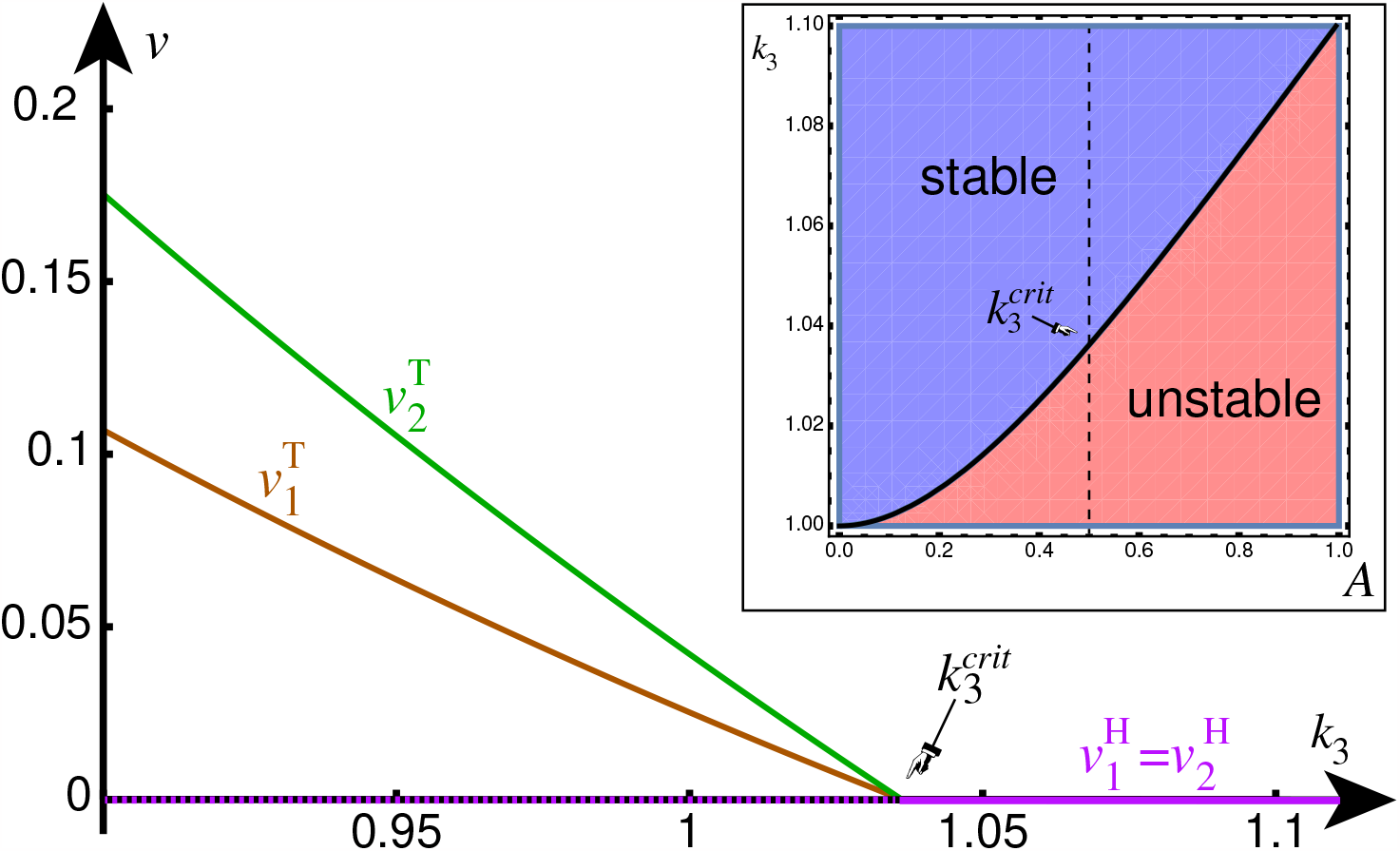
Bifurcation diagram for toxic load in nodes 1 and 2 as a function of toxic clearance *k*_3_; other parameters are *A* = 1*/*2, *ℓ* = 1, *k*_0_ = 1, *k*_1_ = 1, *k*_2_ = 1. Inset: Bifurcation in (*A,k*_3_) parameter space. Increasing activity destabilizes the healthy fixed point by shifting the transcritical bifurcation.

It is interesting to understand the behavior of the toxic equilibrium as a function of the activity. Assuming that activity *A* is small compared to *ℓ*, we can expand the non-trivial equilibrium to first order in *A* to obtain

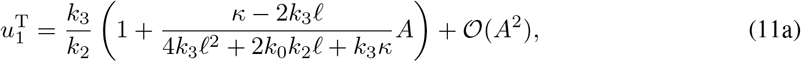

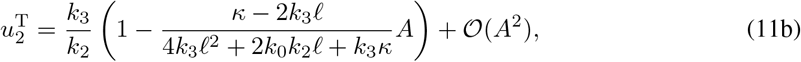

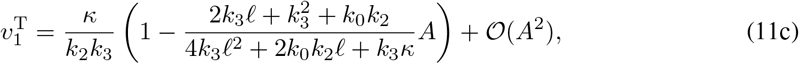

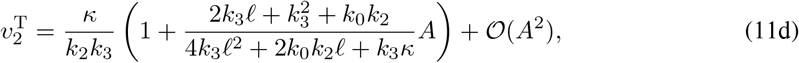

We see that activity can affect the fixed point in two distinct ways assuming *κ >* 0 so that the healthy fixed point is unstable: If *κ*−2*k*_3_*ℓ >* 0, then *u*_1_ increases while *v*_1_ decreases and *u*_2_ decreases while *v*_2_ increases. By contrast, if *κ* −2*k*_3_*ℓ <* 0 is small, then *u*_1_ decreases while *v*_1_ decreases, and *u*_2_ increases while *v*_2_ increases. In the former case, the conversion process is dominating, and the effective conversion at node 1 has decreased while it has increased in node 2. In the latter case, the transport process dominates and both species at node 1 are being shunted over to node 2. This *shunting* phenomenon does not occur in the original heterodimer model and is showcased in Figure 3.

**Figure 3:**
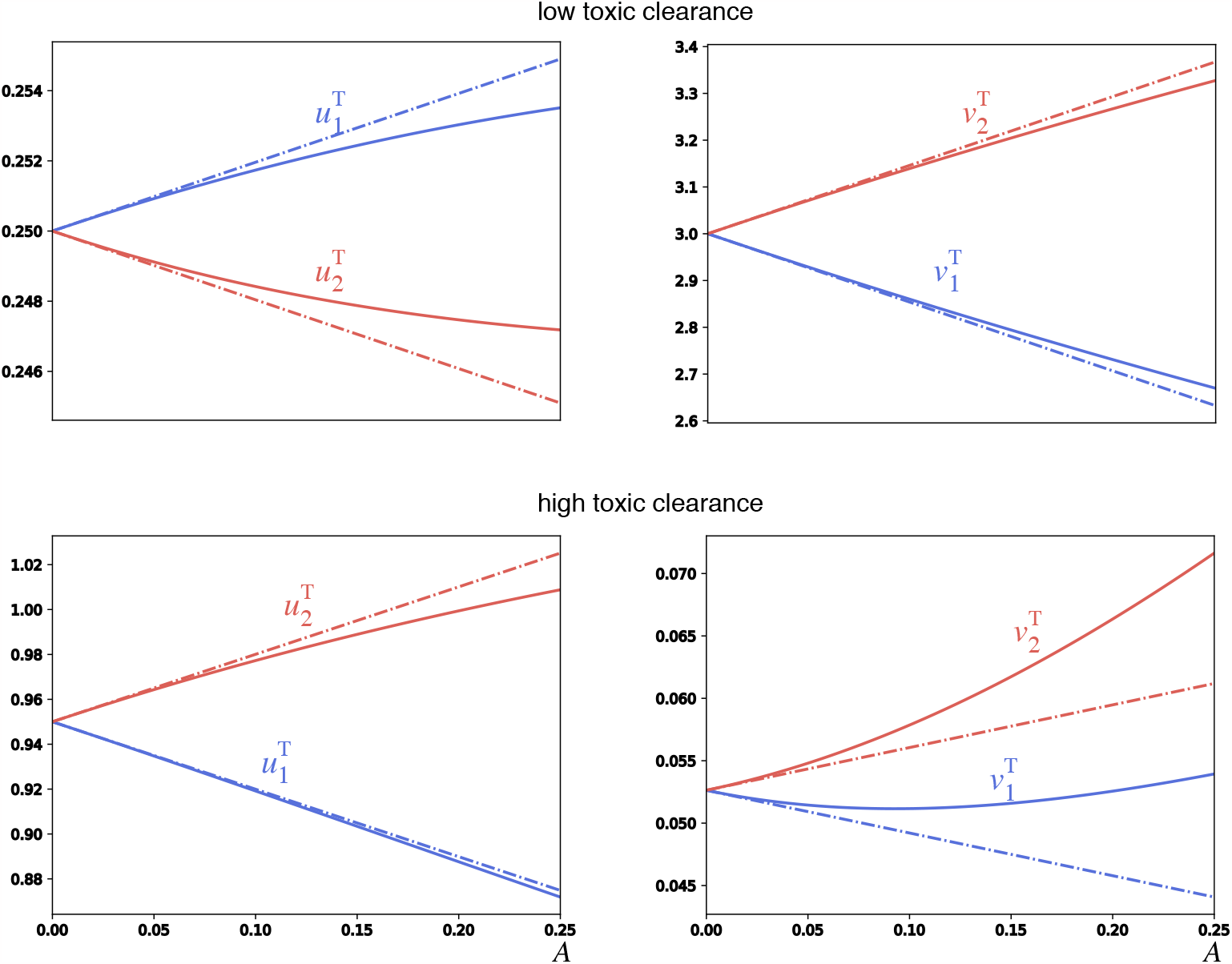
Comparison of steady-states of healthy and toxic species in nodes 1 (blue) and 2 (red) determined by simulation (solid lines) and first-order Taylor expansion (stippled line) of the activity parameter *A*. The upper row demonstrates the conversion-dominated regime, whereas the bottom row demonstrates the transport-dominated regime. All parameters are set to 1, except for *k*_3_ = 0.25 in the first row (far from the transcritical bifurcation at 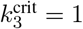) and *k*_3_ = 0.95 in the second row (close to the transcritical bifurcation).

## 3 Coupling heterodimer dynamics with oscillatory activity

In the previous analysis, we considered the activity *A* to be a constant. In the brain, activity may relate to collective neural oscillations that are fast compared to the disease progression. Therefore, we now assume that *A* is determined by the evolution of a pair of phase oscillators with Kuramoto coupling. Since the spreading and activity processes evolve on different time scales, the coupling between the two systems defines a slow-fast dynamical system.

### 3.1 Two coupled phase oscillators

First, consider two phase oscillators, one on each node, with Kuramoto coupling. That is, the state of the oscillation on node *i* ∈ {1, 2} is given by a phase *θ*_*i*_ ∈ 𝕊:= ℝ */*2*π* ℤ that evolves according to

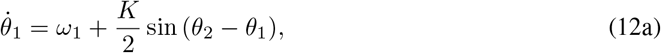

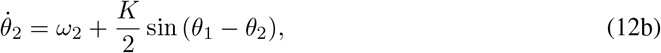

where *ω*_*i*_ *>* 0 are the intrinsic frequencies of the nodes and *K*≥ 0 is the *coupling strength*. Since the coupling depends solely on the phase difference, the dynamics are completely determined by the evolution of the phase difference *ϕ* := *θ*_1_ − *θ*_2_ determined by

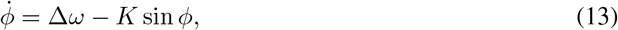

where Δ*ω* = *ω*_1_ − *ω*_2_ is assumed to be positive, without loss of generality. For *K >* | Δ*ω*|there are two fixed points (one unstable and one stable attracting all initial conditions except the unstable fixed point). For *K <* |Δ*ω*|there are no fixed points and any solution *ϕ*(*t*) is periodic. At the critical coupling strength *K* = |Δ*ω*|, there is a saddle-node bifurcation. Hence, we essentially have two regimes depending on the dynamics; we refer to them as the strong-coupling regime (fixed points) and the weak-coupling regime (periodic orbit).

We assume that the activity at each node is related to the *instantaneous frequencies*, 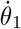 and 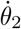of each node. We define the *average frequency* of each oscillator:

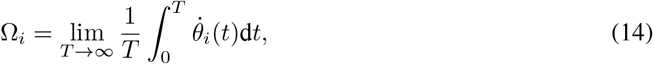

which is independent of the initial conditions. In the strong-coupling regime (the coupling between oscillators is strong compared to the frequency mismatch and the phase difference *ϕ* converges to a fixed point), the oscillators are frequency locked. At the fixed points, we have a constant instantaneous frequency 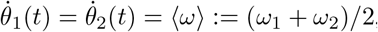, which implies

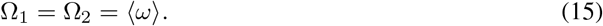

In the weak-coupling regime (the coupling between the oscillators is weak compared to their frequency mismatch and the phase difference undergoes periodic oscillations), we compute the average frequencies Ω_*i*_ through the average frequency difference

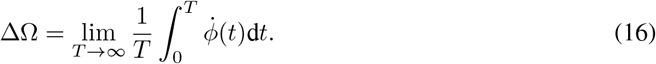

Define Δ*T* = 2*π/*ΔΩ, assume ≤*K <* Δ*ω*, and let *ϕ*(*t*) be the Δ*T* -periodic solution of (13). Note that the sign of 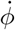 is constant. With (13) we have

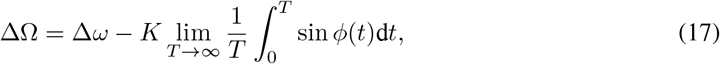

With *m* = *T/*|Δ*T* |, we can rewrite the integral as

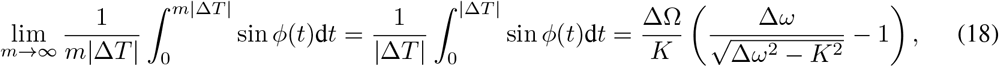

where the last equality follows from substituting *t* by *ϕ* (which is possible since the sign of 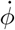 is constant) and solving the resulting integral by Weierstrass substitution. Using this last expression in (17) yields

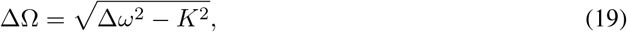

from which we compute the asymptotic frequencies of each node

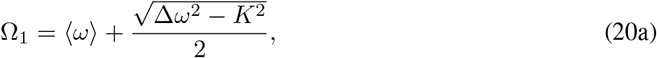

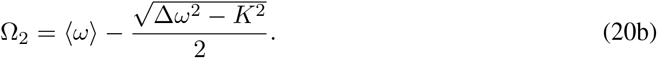

### 3.2 Slow-fast heterodimer-oscillator dynamics

Next, we couple the oscillatory dynamics with the heterodimer model of protein spreading. The two processes will evolve on distinct time scales, determined by a small strictly positive constant *ϵ* ≪1, representing the ratio between the fast activity time scale and the slow spreading time scale. Specifically, the two-node *heterodimer-oscillator* (see Fig. 1c) system is

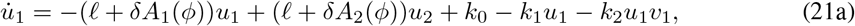

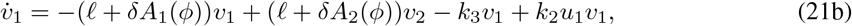

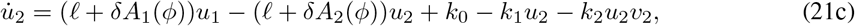

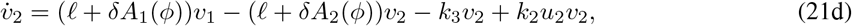

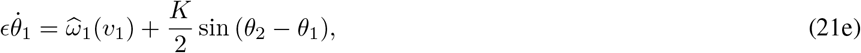

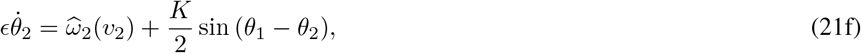

where *δ >* 0 scales the oscillators’ effect on spreading. We assume that the coupling between heterodimer and oscillatory dynamics is through the phase-dependent activity of nodes 1 and 2, that is,

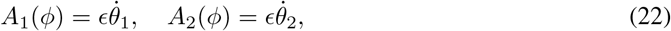

and the intrinsic frequencies

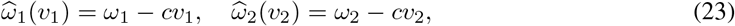

that are decreased by the presence of toxic proteins with a scaling parameter *c >* 0. As discussed above, we may replace the phase dynamics in (21) by the evolution of the phase difference *ϕ* as above given by

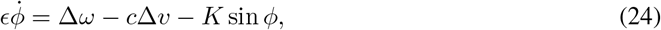

where Δ*v* = *v*_1_ − *v*_2_ is the difference in toxic protein concentration. The phase locking behavior is now determined by the effective intrinsic frequency difference 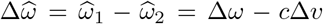, which is a function of Δ*v*. As there is no sensible interpretation of *negative* neuronal activity, we will only consider parameters for which *A*_i_ (*t*) ≥ 0, *i* ∈ {1, 2} for all *t*.

Given that the spreading dynamics is much slower than the oscillator dynamics (on the order of years versus seconds), we are interested in the dynamics for small *ϵ* close to the singular limit *ϵ*→0. In the singular limit, the phase dynamics relax instantaneously to the asymptotic dynamics of the phase-difference *ϕ*(*t*). Thus, the dynamics in the singular limit depend on which dynamical regime the phase difference is operating in. In the phase-locked regime, the dynamics relax instantaneously to equilibrium, which defines the critical manifold of the slow-fast system on which *A*_*i*_ takes its value at equilibrium. In the regime where the phase difference *ϕ*(*t*) is drifting, we replace the instantaneous frequency in *A*_*i*_ by the temporal average Ω_*i*_; this is similar to the approach in [15]. Finally, we consider the system at the border between the two regimes.

### 3.3 The phase-locking regime

Assume that 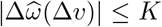. Then the singular-limit dynamics on the slow manifold is determined by the stable phase-difference equilibria

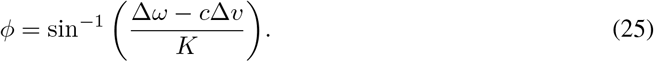

Inserting the fixed point into *A*_*i*_, both nodes have identical activities

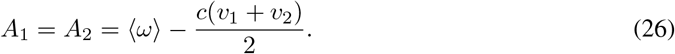

Substituting these expressions into the slow system gives us the dynamics on the phase-locking critical manifold.

Since the nodes have identical activity levels, the dynamics are qualitatively equivalent to those of the isolated heterodimer model. In particular, the system has the same pair of healthy and toxic fixed points as the heterodimer model given by (4) :

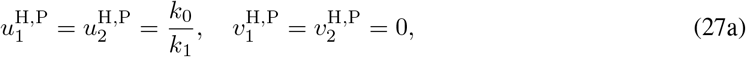

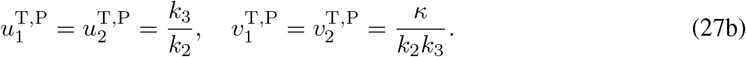

The stability of the healthy fixed point is determined by the eigenvalues of the Jacobian matrix:

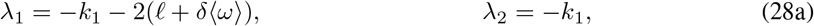

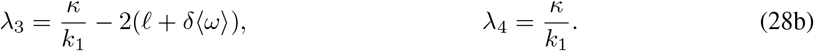

Thus, the healthy fixed point loses its stability at *κ* = 0.

The assumption of being in the phase-locking regime, 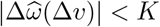, gives a consistency condition for the existence of the fixed points on the critical manifold. Note that since the activity of each node in the phase-locking regime is identical, the slow dynamics is symmetric in the sense that exchanging the two nodes has no impact on the dynamics. Furthermore, for both fixed points, the nodes are “equal” in the sense that they take the same state and satisfy Δ*v* = 0. Thus, the healthy and toxic fixed points only exist as fixed points on the critical manifold for the slow dynamics if 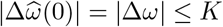.

### 3.4 The drifting regime

Outside the phase-locked regime, 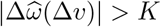, the fast oscillatory dynamics do not relax to equilibrium but evolve on a periodic orbit. As these oscillations are much faster than the evolution of the slow dynamics, we average out the fast oscillations by replacing the activities *A*_*i*_ by their temporal averages to define the *drifting regime*. Specifically, replacing *ω*_*i*_ with 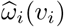 in (20) and assuming, without loss of generality, that 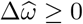 yields (20) the activities

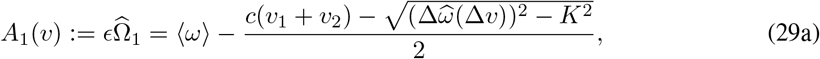

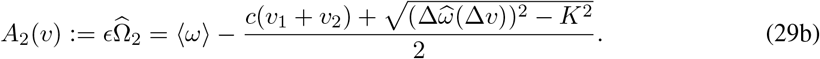

Substituting these activities into the dynamical equations for the slowly-evolving heterodimer equations yields the dynamics of the drifting regime. As the activities of the two nodes are now distinct, the dynamics is similar to the skewed heterodimer model in Section 2.2. There is one healthy fixed point *u*^H,D^ in the drifting regime with coefficients

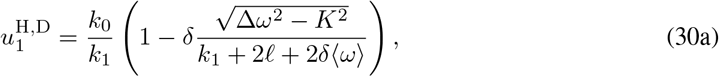

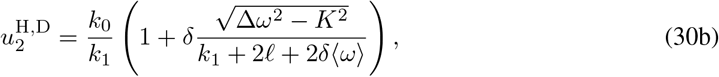

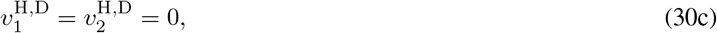

under the assumption that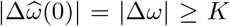. Similar to the skewed heterodimer model, the symmetry of the fixed points is broken. Linear stability of the healthy fixed point is determined by the eigenvalues of the Jacobian matrix:

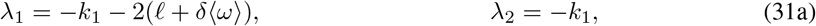

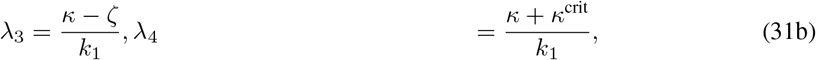

where

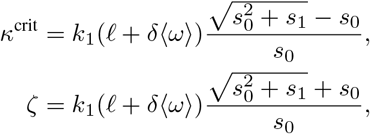

with *s*_0_ = 2*k*_1_ (*ℓ* + *δ* ⟨*ω*⟩) (*k*_1_+2*ℓ*+2*δ* ⟨*ω*⟩) and *s*_1_ = 4*δ*^2^*k*_0_*k*_2_(Δ*ω*^2^−*K*^2^)(*k*_0_*k*_2_+*k*_1_ (*k*_1_ + 2*ℓ* + 2*δ* ⟨*ω*⟩)). Remembering that the healthy fixed point only exists for |Δ*ω*| *> K*, we can assert that *κ*^crit^, *ζ >* 0, as is verifiable by inspecting *s*_1_. As such, *λ*_4_ determines the stability of *u*^H,D^. The critical value for 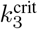 at which the transcritical bifurcation occurs is

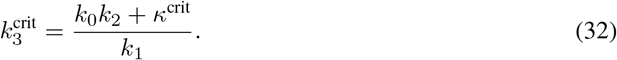

Assuming *δ* small compared to *ℓ*, we expand the toxic fixed point *u*^T,D^ in the drifting regime to first order in *δ*, giving us

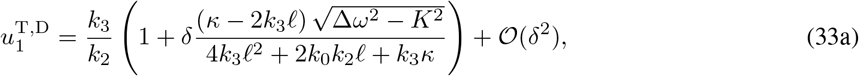

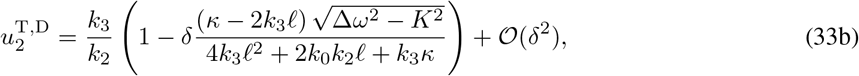

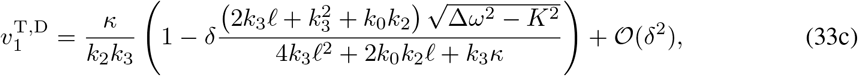

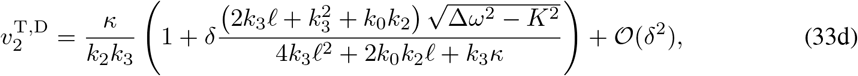

where the coefficients are similar to the expansion of the skewed heterodimer toxic fixed point, except that they are scaled by 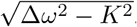. As such, we have transport- and conversion-dominated behavior for small and large values of *κ*−2*k*_3_*ℓ* respectively. More importantly, we have established the existence of a toxic fixed point *u*^T,D^ on the drifting regime for small *δ*.

Note that the above coefficients are only defined for *K* ≤ |Δ*ω*|, similarly to *u*^H,D^. As such, our preceding analysis suggests a symmetry-breaking, global bifurcation occurring at *K* = |Δ*ω*| in which from one side (from the phase-locking regime) two fixed point branches collide and disappear (saddle-node bifurcation on an invariant circle), but from the other side (from the drifting regime) two periodic solutions collide and disappear. Furthermore, the fixed points in the phase-locking regime are symmetric between the nodes with respect to their heterodimer variables, whereas both the periodic solutions are asymmetric in this respect. A summary of the heterodimer-oscillator dynamics in the strong-coupling and weak-coupling regimes can be found in Figure 4 alongside numerical solutions for *ϵ >* 0. Moreover, an overview of the dynamical regimes and the (singular-limit) transcritical bifurcation is illustrated in (*K, k*_3_) parameter space in Figure 5.

**Figure 4:**
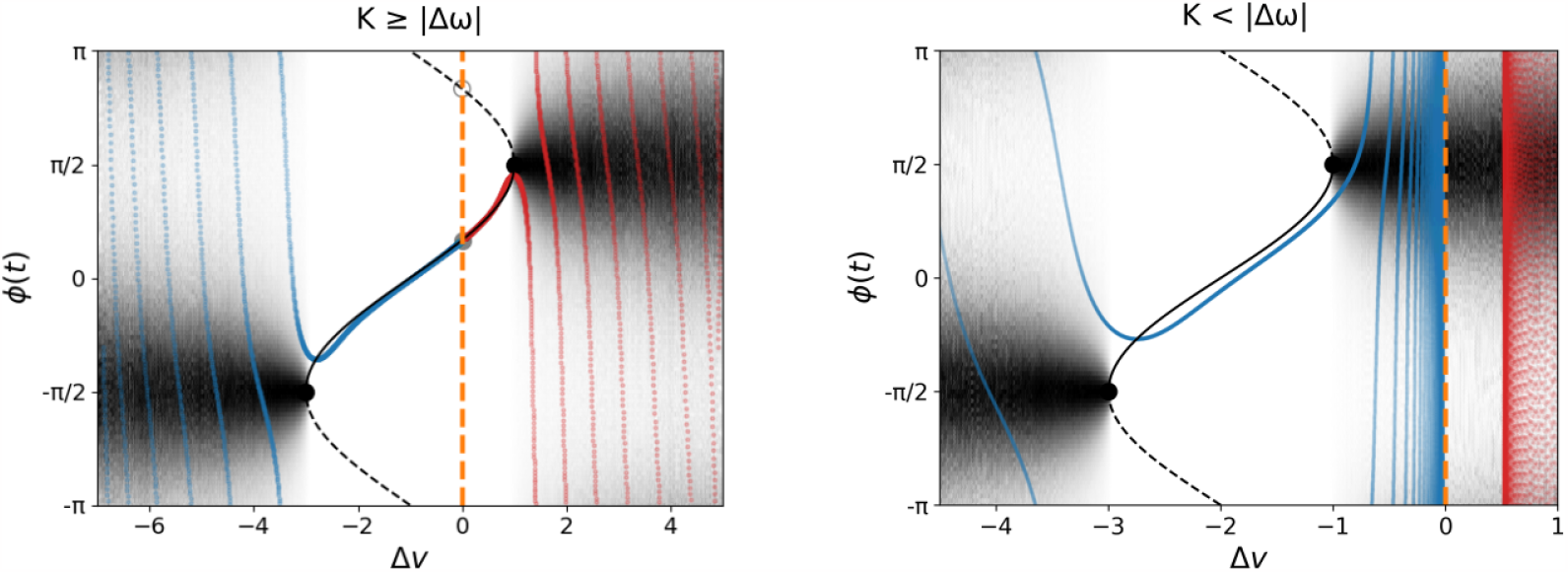
Summary of the dynamics in the phase-locking and drifting regime with simulations in the healthy (blue) and toxic (red) regimes of the heterodimer-oscillator with *ϵ >* 0. *Left:* Summary for *K >*Δ*ω* over the phase-difference and toxic species difference, where *K* = 2, Δ*ω* = 1, *c* = 1, *k*_0_ = 1, *k*_1_ = 1, *k*_2_ = 1, *δ* = 1, *ℓ* = 10^−3^ with forward solutions in the toxic (*ϵ* = 0.2, *k*_3_ = 0.75) and healthy regime (*ϵ* = 0.075, *k*_3_ = 1.25). Both forward solutions are symmetric with respect to the slow variables. *Right:* Summary for *K <* Δ*ω* where *K* = 1, Δ*ω* = 2, *c* = 1, *k*_0_ = 1.5, *k*_1_ = 1, *k*_2_ = 1, *δ* = 1, *ℓ* = 10^*−*3^ and with forward solutions in toxic (*ϵ* = 0.1, *k*_3_ = 0.125) and healthy regimes (*ϵ* = 0.075, *k*_3_ = 1.25). Both forward solutions are asymmetric with respect to the slow variables (the healthy solution is asymmetric with respect to the healthy species). Note that the trajectories in the healthy regimes converge to Δ*v* = 0 (highlighted with a stippled, orange line) in both diagrams.

**Figure 5:**
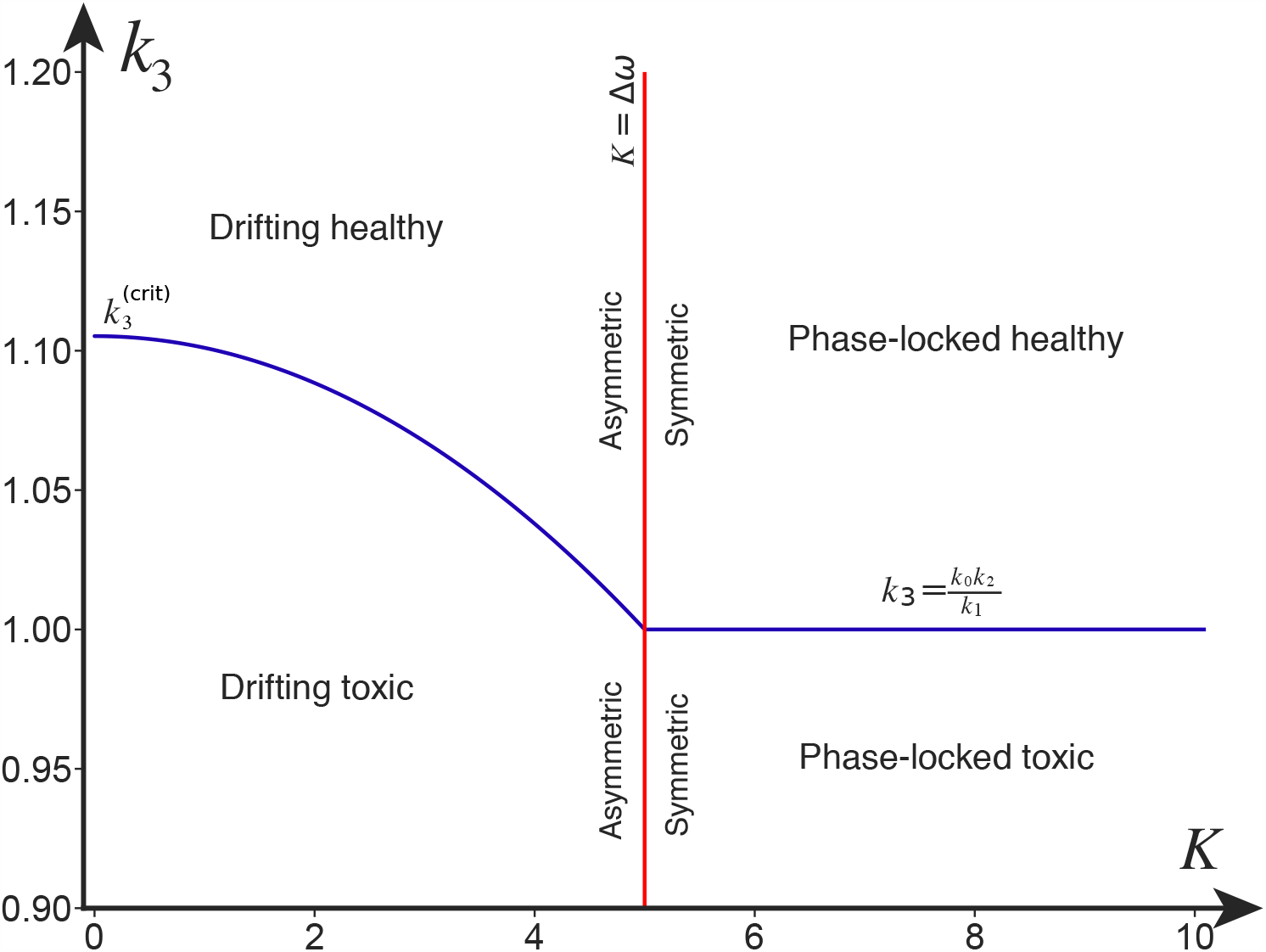
Summary of the dynamics of the 2-node heterodimer-oscillator in the singular limit (*ϵ*→ 0). The labels in each quadrant state which fixed point we know to be stable. The transcritical bifurcation in the weak-coupling and strong-coupling regime is presented, together with the breaking of the symmetry between the two nodes in the fixed points, which occurs at *K* = |Δ*ω*|. Parameters are *k*_0_ = 1, *k*_1_ = 1, *k*_2_ = 1, *ℓ* = 1, |*ω*_1_| = 10, *ω*_2_ = 5, *δ* = 5.

### 3.5 Transitions between the phase-locking and drifting regimes

With an understanding of the dynamics within the phase-locking regime (Section 3.3) and drifting regime (Section 3.4) at hand, we can now elucidate possible transitions between the regimes. The boundary between the regimes is where the fast dynamics undergo a saddle-node bifurcation at 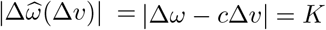. Equivalently, we obtain the following condition for the regime border

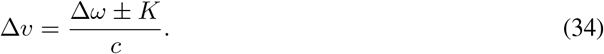

The value of Δ*v* is subject to the slow dynamics (21). Specifically, the sign of 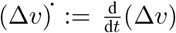 termines the transitions between the phase-locking and drifting regimes: For the right boundary of the phase-locking regime, 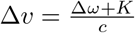, negative (Δ*v*)^**·**^ indicates that the slow flow points from the drifting regime into the phase-locking regime and a positive (Δ*v*)^**·**^in the opposite direction. For the left boundary, the conditions are the other way around. In the following, we will argue that, under certain assumptions, the flow points towards the phase-locking regime for *K >* |Δ*ω*| and towards the drifting regime for *K <* |Δ*ω*| ; this is sketched in Figure 6.

**Figure 6:**
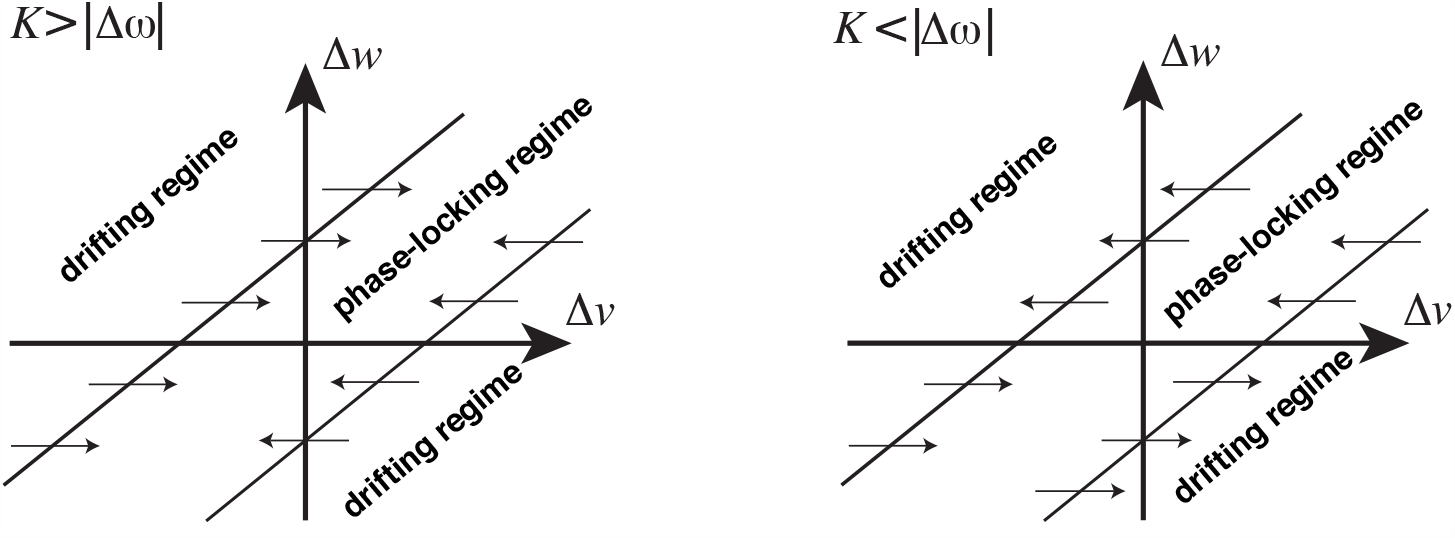
The vector field of Δ*v* in terms of Δ*ω* and Δ*v* in the strong-coupling (left) and weak-coupling regime (right). The inner region in both diagrams is the phase-locking regime, and the outer regions are the drifting regime. We see that for strong coupling *K >* |Δ*ω*|, the vector field points inwards to the phase-locking regime. However, for the weak-coupling regime *K <* |Δ*ω*|, the vector field points to the left-hand drifting regime for Δ*ω >* 0 (node 1 is more active than node 2) and to the right-hand drifting regime for Δ*ω <* 0 (node 2 is more active than node 1).

To determine the transitions between the regimes, we consider the dynamics of Δ*v*. In the singular limit, the dynamics have relaxed to the saddle-node equilibrium and thus *A*^*∗*^ := *A*_1_ = *A*_2_. Now we assume that the *u*_*i*_ take their equilibrium values, with *u*_1_ = *u*_2_ = *k*_0_*/k*_1_ (healthy regime) and *u*_1_ = *u*_2_ = *k*_3_*/k*_2_ (toxic regime); *u*_1_ = *u*_2_ =: *u*^*∗*^ in either case. According to (21), the evolution of Δ*v* is determined by

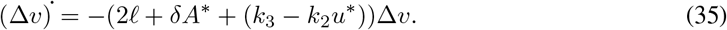

We claim that the first factor is not positive (i.e., the quantity in the parentheses is not negative). The first two terms are clearly positive since *ℓ* ≥ 0 and, by assumption, *A*^*∗*^ ≥ 0. For the third term, *k*_3_−*k*_2_*u*^*∗*^ ≥ 0 is equivalent to 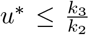. But, by assumption, *u*^*∗*^ = *k*_3_*/k*_2_ or *u*^*∗*^ = *k*_0_*/k*_1_ ≤ *k*_3_*/k*_2_ so in either case the third term is not negative. We conclude that the sign of (Δ*v*)^**·**^ only depends on the sign of Δ*v*. For the right boundary of the phase-locking regime, we have Δ*v* = (Δ*ω* + *K*)*/c* so Δ*v >* 0 or equivalently *K >*−Δ*ω* implies (Δ*v*)^**·**^ ≤ 0 (flow towards the phase-locking regime). Conversely, *K*−*<* Δ*ω* implies (Δ*v*)^**·**^ ≥ 0 (flow towards the drifting regime). Similarly, for the left boundary of the phase-locking regime, we have Δ*v* = (Δ*ω*−*K*)*/c* so *K <* Δ*ω* implies (Δ*v*)^**·**^ ≤ 0 (flow towards the drifting regime) and *K >* Δ*ω* implies (Δ*v*)^**·**^ ≥0 (flow towards the phase-locking regime).

Thus in terms of the system parameters, the crucial quantity is the oscillator coupling relative to the intrinsic frequency mismatch. If *K >* |Δ*ω*| then the flow points towards the phase-locking regime on either boundary. If *K <* |Δ*ω*| then the flow points in the same direction on each boundary and the direction is determined by the sign of Δ*v*. These cases are illustrated in Figure 6.

### 3.6 Extending the parameter regime

From the beginning, we have assumed *c* and *δ* to be positive. These assumptions, however, may not be fit for all applications of the heterodimer-oscillator model. For example, one might envision spreading processes that *increases* oscillatory activity locally (*c <* 0), and, in return, oscillatory processes that *decreases* spreading to its neighboring oscillators (*δ <* 0). First, we consider the case *c <* 0. None of the singular-limit fixed points nor their stability depend on *c*, and the regime border analysis above can be repeated successfully for *c <* 0 and *δ >* 0 (noting that the left- and right-hand borders swap places). For *δ <* 0, we may assume *δ* ⟨*ω*⟩ *> − ℓ* to guarantee that our stability analysis of the phase-locking and drifting equilibria remains unaffected (see eigenvalues in Eqs. (28) and (31)). The assumption is within reason; it is equivalent to stating that the link between the nodes *ℓ* + *δA*(*ϕ*) does not change signs for Δ*v* = 0. For the regime border analysis to hold, we require 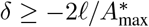 where 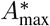is the maximum of the phase-locked activity over *v*_1_ and *v*_2_ (see Eq. (26)). For *c >* 0, we have that 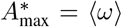 giving *δ* ⟨*ω*⟩≥−2*ℓ*, which is already satisfied by ⟨*δ ω*⟩*>*−*ℓ*. However, 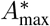 increases indefinitely in *v*_1_ and *v*_2_ for *c <* 0. Hence, we need additional bounds on the variables *v*_1_ and *v*_2_ to ensure that the regime border analysis holds. Although the regime border analysis cannot be repeated for *c, δ <* 0 without further assumptions, we can conclude that the fixed point linear stability analysis generalizes to *c, δ* ∈ ℝ,

## 4 Activity-spreading feedback on networks

Investigating the dynamics of the heterodimer-oscillator system on more general networks, we find that the results from the 2-node heterodimer-oscillator system provide a strong intuition for the generalized network dynamics. Specifically, we consider a network of *N* nodes determined by the *N*×*N* (weighted) adjacency matrix **W** with Laplacian **L**. Let *u*, ∈ *v* ℝ ^*N*^ denote the healthy and toxic species concentration at each node and *θ* ∈ 𝕊 ^*N*^ the state of the oscillators on each node. Generalizing (21), the states evolve according to

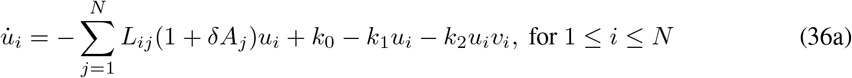

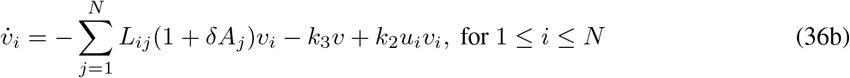

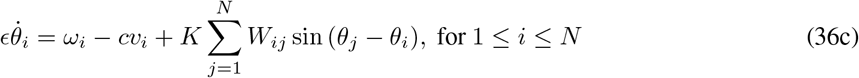

where 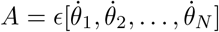.

### 4.1 Numerical exploration of key example networks

#### Erdős–Rényi random graphs

Dynamics on Erdős–Rényi random graphs retain the transcritical bifurcation near *κ* = 0 alongside its symmetry for small differences between the intrinsic frequencies of the nodes; cf. Figure 7. However, with large differences in the intrinsic frequencies, the transcritical bifurcation extends the toxic parameter regime and breaks the symmetry of the fixed points between the nodes, just as expected from our analysis of the 2-node system.

**Figure 7:**
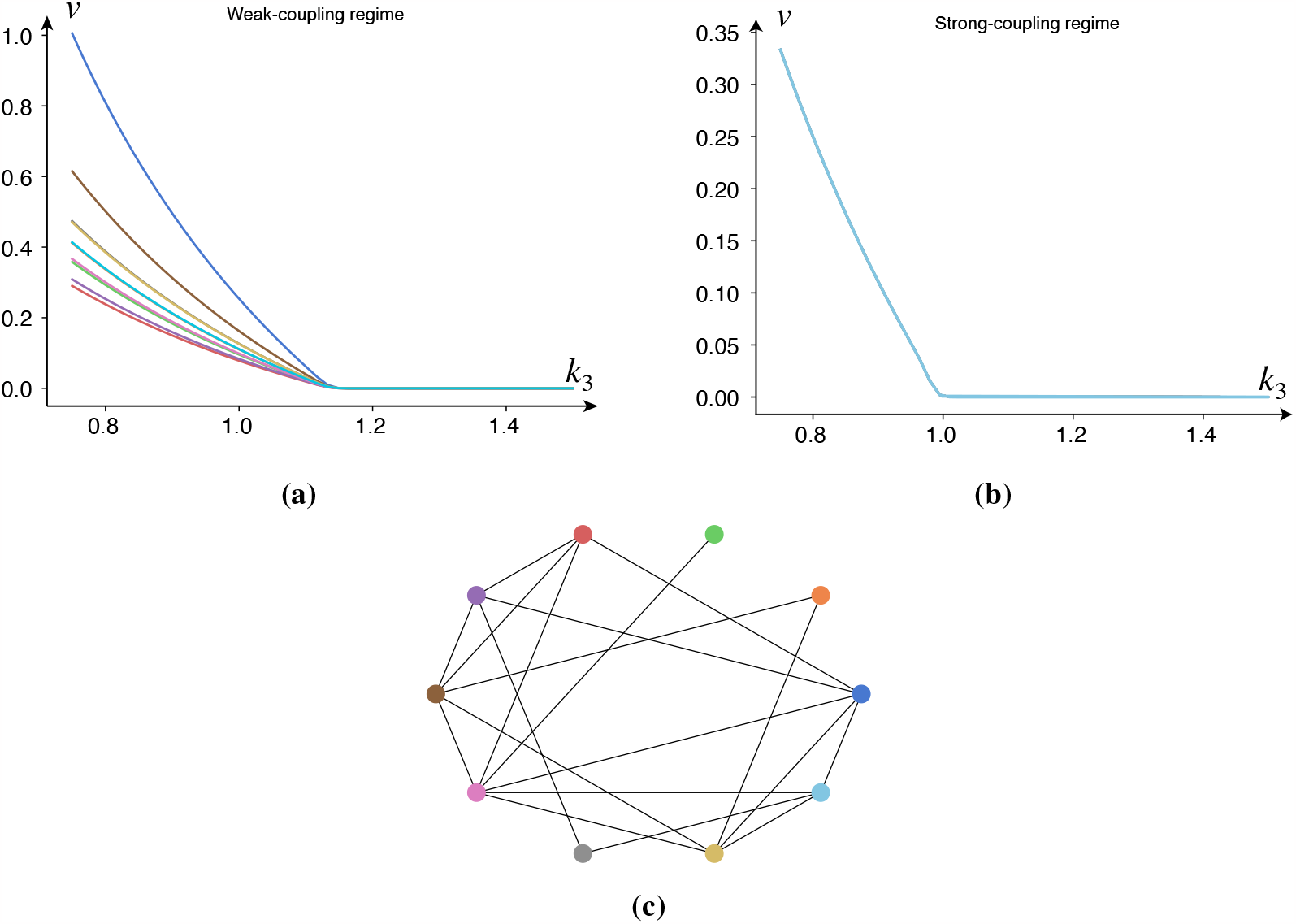
Simulations demonstrating the transcritical bifurcation during the weak-coupling and strong-coupling regime in a random graph. All weights in the network are set to 1, and the intrinsic frequencies were drawn from a normal distribution. *(a)* The weak-coupling parameters are *ρ* = 0.1, *k*_0_ = 1, *k*_1_ =1, *k*_2_ = 1, *E*(*ω*) = 10, Var(*ω*) = 0, *c* = 0.5, *ϵ* = 10^−3^, *δ* = 10, *K* = 0.1, while one of the oscillators (dark blue) at a slower frequency *ω* = 5. *(a)* The strong-coupling parameters are *ρ* = 0.1, *k*_0_ = 1, *k*_1_ = 1, *k*_2_ = 1, *E*(*ω*) = 10, Var(*ω*) = 0.25, *c* = 0.5, *ϵ* = 10^−3^, *δ* = 10, *K* = 0.1. *(c)* The Erdős–Rényi graph (*N* = 10, *p* = 0.5).

#### Chain graphs

To further test our intuition, we create a chain graph with decreasing frequencies along the chain. If we initialize a small amount of toxic species in each node, we expect (in the steady state) a gradient of increasing toxic species along the chain. As observed in Figure 8, this prediction is accurate. Additionally, we observe shunting behavior. First, the healthy species are quickly transported according to the nodes’ activity gradient, and then the healthy species are converted into toxic species. According to our 2-node analysis, such shunting behavior should occur close to the original transcritical bifurcation *κ* = 0, which is where the simulation in Figure 8 has been parameterized.

**Figure 8:**
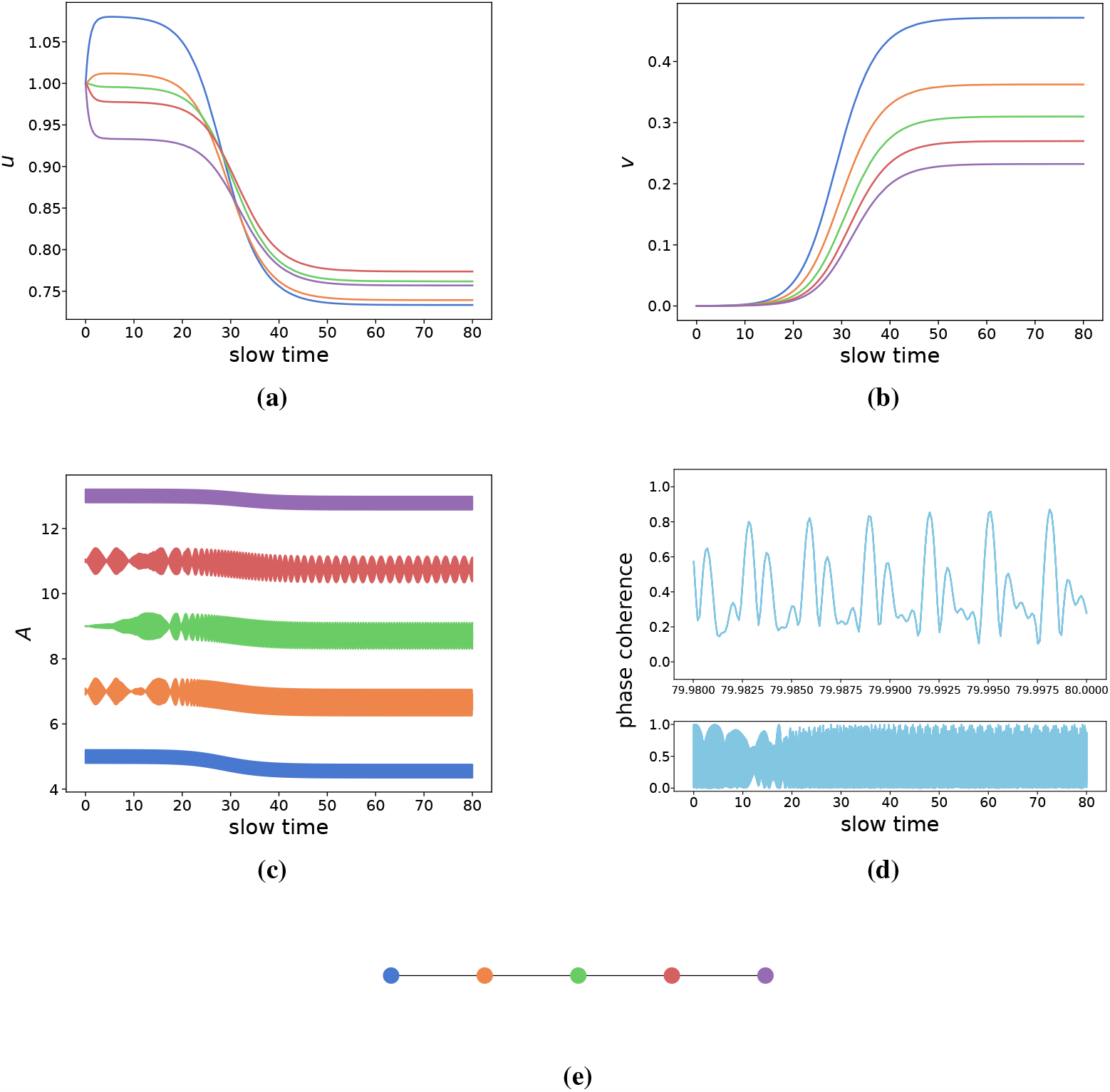
Simulations demonstrating heterodimer-oscillator dynamics on a chain network. All link weights are set to 0.1 while other parameters are *ρ* = 0.5, *k*_0_ = 1, *k*_1_ = 1, *k*_3_ = 0.75, *k*_2_ = 1, *ϵ* = 10^−3^, *δ* = 1 *K* = 1, *c* = 0.5. The natural frequencies of the nodes, *ω*_*i*_, range from 5 to 15 with increments of 2.5. Colors are consistent across figure panels. *(a)* The evolution of healthy species. *(b)* The evolution of toxic species. *(c)* The activity (instantaneous frequencies) of the nodes. *(d)* The phase-coherence of the Kuramoto order parameter of the oscillators. *(e)* Graph of the network.

#### Clustered networks

Many complex networks show high degrees of clustering. As such, we created a network of 3 fully-connected clusters of 10 nodes each, where each cluster is connected to each other by two links chosen between a random node pair; cf. Figure 9. We then drew the intrinsic frequencies from normal distributions where each cluster has a different mean. That is, one cluster will be highly active, one will be moderately active, and one will be less active. By doing so, we will have 3 synchronized clusters that are weakly connected to each other. As before, we set the parameters in the toxic regime, yet close to the original transcritical bifurcation at *κ* = 0. As shown in Figure 9, the simulations confirm the intuition from the 2-node system. At first, the healthy species are shunted towards the lesser active clusters, where they are subsequently converted into toxic species. The least active cluster thus produces the most toxic species followed by the moderately active and highly active clusters, respectively. These simulations suggest that the heterodimer-oscillator might also be suitable for mean-field models of population dynamics.

**Figure 9:**
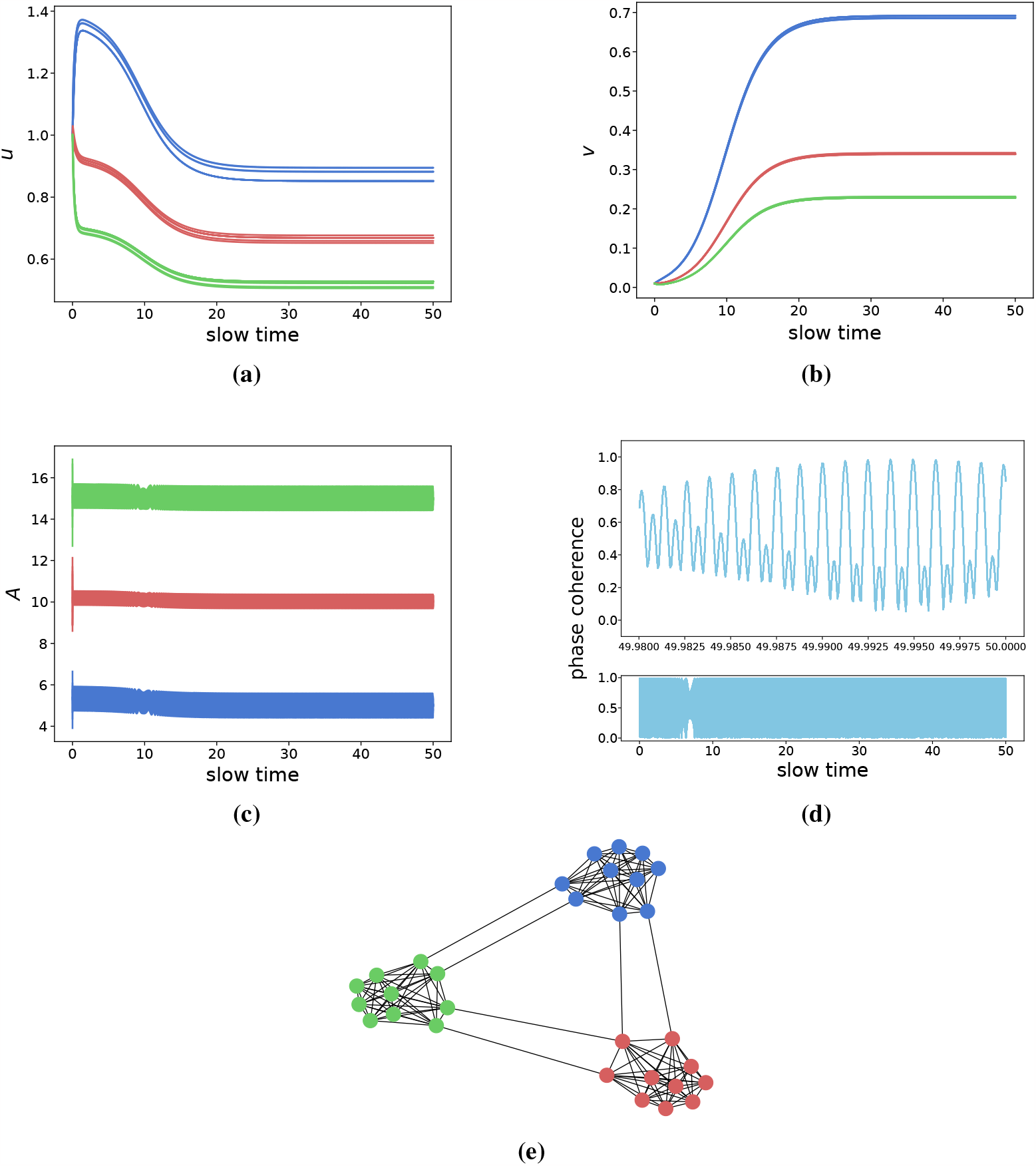
Simulations demonstrating heterodimer-oscillator dynamics on a clustered network. All network link weights are set to 0.1, while other parameters are *ρ* = 0.5, *k*_0_ = 1, *k*_1_ = 1, *k*_3_ = 0.75, *k*_2_ = 1, *ϵ* = 10^−3^, *δ* = 1 *K* = 1, *c* = 0.5. Clusters are colored green (highest activity), red (medium activity), and red (lowest activity) respectively. The average intrinsic frequencies of the clusters, *ω*_*i*_, are normally distributed with means 5 (blue), 10 (red), and 15 (green) and a common standard deviation of 0.5. *(a)* The evolution of healthy species. *(b)* The evolution of toxic species. *(c)* The activity (instantaneous frequencies) of the nodes *(d)* The phase-coherence of the Kuramoto order parameter of the oscillators. *(e)* Graph of the network.

### 4.2 Exploring coevolutionary dynamics in Alzheimer’s disease

Our original motivation was to investigate the effect that the slow-fast dynamics between neuronal activity and pathological protein spreading exert on the progression of neurodegenerative diseases. Previous studies have modeled the progression of Alzheimer’s disease as a spreading process on network reconstructions of the human brain. These network reconstructions are called connectomes and are built using DTI imaging which are subsequently parcellated into networks of arbitrary size. Nodes in the network represent brain regions and links between brain regions represent axonal bundles.

Typically, modelers will simulate the spreading of tau proteins across the connectome, leading to neurodegeneration and neuronal death. Tau proteins start to aggregate at the entorhinal cortex and spread progressively to the hippocampus, the limbic system, and the neocortex. The successive spread of tau has been shown to follow a pattern, and, as such, the spreading of tau is divided into six stages known as the Braak staging scheme. However, not all patients follow the Braak staging scheme. In fact, studies suggest that Alzheimer’s patients fit into different subgroups based on their staging patterns [24, 25]. Furthermore, as noted in the Introduction, tau proteins are believed to be transported at a higher rate from higher-active neurons [26], and several studies suggest a crucial link between brain-wide correlations of brain activity and disease spreading patterns [42, 43]. We here provide proof-of-concept, with our heterodimer-oscillator model, that neuronal activity may play a mechanistic role in the spreading of tau protein seen in Alzheimer’s disease staging.

We simulate the spreading of tau on the 83-node Budapest Reference Connectome [44]—in which the simulation initially follows the canonical Braak staging pattern—and gradually increase the effect that neuronal activity has on spreading, which is achieved by increasing *δ*. To simulate the natural progression of Alzheimer’s disease, we only initialize a nonzero concentration of toxic protein in the entorhinal cortex (Braak stage I). As illustrated in Fig. 10(a,b), we find that increasing the effect that neuronal activity asserts on spreading dynamics changes the staging pattern of tau, and also breaks the symmetry of toxic tau concentration—which is inherited by the original oscillator-less heterodimer model—suggesting that the inclusion of neuronal activity may benefit our understanding of the variable severity of tau pathology in different brain regions. Furthermore, we find that changing *δ* (intensifying the feedback loop) has little effect on the asymptotic oscillator dynamics (asymptotic as in the average dynamics close to the end of the spreading simulation), as seen in Fig. 10(c). Note that the toxic proteins *do* slow down the frequency of the oscillators, however, the amount of slowing does not appear to depend significantly on *δ*. Likewise, the average phase-coherence also appears unfazed by changing *δ*, as shown in Fig. 10(d).

**Figure 10:**
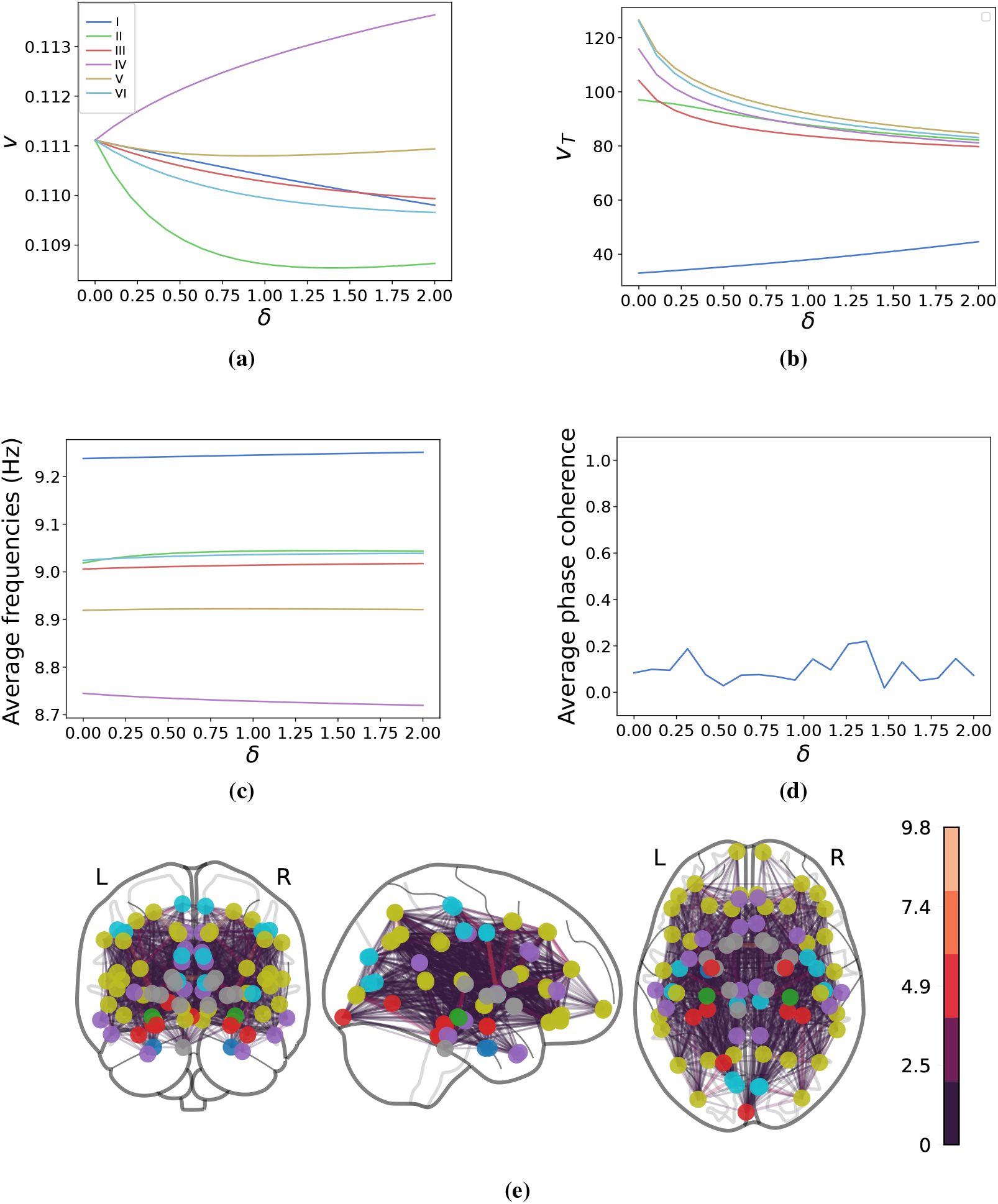
Simulations of toxic tau spreading across an 83-node human connectome showing the asymptotic (steady-state) behavior of the system as a function of *δ* (the effect activity has on spreading). Parameters are *ρ* = 0.001, *k*_0_ = 1, *k*_1_ = 1, *k*_3_ = 0.9, *k*_2_ = 1, *ϵ* = 0.01, *K* = 0.1, *c* = 10. The natural frequencies, *ω*_*i*_, are drawn from a normal distribution with a mean 10 and standard deviation of 0.5. Nodes are initialized with *u*_*i*_ = 1 and *v*_*i*_ = 0, apart from the entorhinal cortices which are initialized with *u*_*i*_ = 1 and *v*_*i*_ = 0.1 (nonzero toxic concentration). *(a)* Average asymptotic amount of tau species in each Braak stage. *(b)* Average time for regions in different Braak stages to become infected with tau. *(c)* Asymptotic average frequencies of the oscillators at the end of the slow-time simulation. *(d)* Average global phase-coherence over the entire slow-time spreading simulation. *(e)* Graph of the brain network with edges colored according to their weight. Nodes that are not part of any Braak stage are colored gray.

## 5 Discussion

Reduction to slow manifolds and ad hoc averaging allows us to elucidate the heterodimer-oscillator dynamics in the singular limit. The heterodimer-oscillator operates in two regimes, as the oscillators are either phase-locked or drifting. In the phase-locking regime, the heterodimer-oscillator exhibits dynamics similar to the original heterodimer model, whereas in the drifting regime, the dynamics is similar to the skewed heterodimer model. In both regimes, we identify a pair of healthy and toxic fixed points exchanging stability at a transcritical bifurcation (for small *δ* in the drifting regime). Inspecting the evolution equations for the heterodimer-oscillator, we see that it inherits the symmetric fixed points of the heterodimer model when *A*_1_ = *A*_2_ (under phase-locking), as the transport terms cancel each other out. Additionally, there is a unique healthy fixed point in both the drifting and phase-locking regimes; setting *v*_1_ = *v*_2_ = 0 reduces the equilibrium conditions to a determinate system of linear equations. The healthy fixed point only exists in the phase-locking regime when *K* ≤|Δ*ω*|, and likewise, the healthy fixed point only exists in the drifting regime *K* ≥|Δ*ω*|. However, the nonlinearities introduced by the heterodimer-oscillator may introduce novel (toxic) equilibria beyond those identified herein. We also established—when healthy species *u* are close to their steady-state values—the direction of the vector field of Δ*v* at the border between the phase-locking and drifting regimes. In doing so, we ruled out the possibility of stable limit cycles crossing the regime border at the singular limit. However, we cannot rule out the existence of limit cycles within either of the regimes. Nonetheless, it seems likely that the heterodimer-oscillator approaches the phase-locking regime for *K* ≥|Δ*ω* | and the drifting regime for *K <* |Δ*ω*|.

The large timescale separation one typically has for the evolution of neurodegenerative diseases justifies the analysis of the singular limit. While smaller timescale separation may alter the dynamics of the system, the numerically obtained solution trajectories shown in Fig. 4 suggest that the heterodimer-oscillator is well-approximated by the singular-limit dynamics even for *ϵ* ≈ 0.1. In particular, numerical simulations indicate that the stability of the phase-locking and drifting regimes is accurately described by the singular limit analysis, even for complex networks. However, we cannot rule out the occurrence of more complex dynamical phenomena for larger *ϵ*, though it has not yet been observed.

In addition to a smaller timescale separation, generalizations of the oscillator dynamics and the coupling between oscillator and spreading dynamics are likely to induce new dynamical phenomena. We chose a simple relationship between the heterodimer dynamics and the oscillator intrinsic frequencies by setting 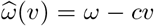, which can be interpreted as a first-order Taylor expansion of a more complicated 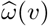. From the expansion of the toxic fixed point in the drifting regime in the singular limit, we have that 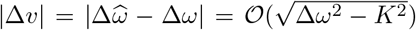 which implies that 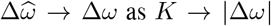. In other words, the heterodimer-oscillator SNIC bifurcation at 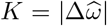 becomes indistinguishable from *K* = |Δ*ω*|. As such, the Kuramoto oscillators appear to only depend on the oscillator parameters. As such, higher-order terms in 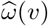 may be necessary to induce qualitative changes in Kuramoto dynamics. A smaller timescale separation may also lead to more interesting behavior in the Kuramoto dynamics. If the temporal changes of the effective intrinsic frequencies 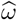 are faster, there may be transient chimera-like states, where parts of the network are severely affected by the spreading and others are not, leading to clusters synchronized at different frequencies. Another interesting scenario involves several heterodimer species on the same network. These species may interact [35] and even affect the underlying oscillator dynamics differently [41]. The latter case could be realized by having two heterodimer species evolving, where one speeds up (negative *c*) and the other slows down (positive *c*) the intrinsic frequencies of the oscillators. It is also possible to modify the heterodimer-oscillator to affect the coupling strengths between oscillators as opposed to their intrinsic frequencies. With the addition of coupling strength adaptation rules, we expect more elaborate oscillator dynamics in the symmetry-breaking (drifting) regime in line with previous research [13].

We have focused on oscillatory processes that accelerate spreading (*δ >* 0) and spreading processes that slow oscillatory processes (*c >* 0). Nonetheless, the heterodimer-oscillator may be fit for modeling phenomena where either *c* or *δ* are negative. By asserting a lower bound on *δ*, we can extend the linear stability results to *c, δ* ∈ ℝ. However, we cannot repeat the regime border analysis when both *c <* 0 and *δ <* 0. In this case, one can imagine a positive feedback loop leading to a stable toxic solution for large Δ*v*, independently of the stability of the healthy fixed point; the node with the highest toxic concentration will increase in activity, causing a reduction in outward transport, followed by an increase in toxic concentration and so on. Still, we are yet to identify applications to which both *c* and *δ* should be negative.

The formulation of the heterodimer-oscillator was primarily motivated by the case of Alzheimer’s disease and other neurodegenerative diseases. The impact of neuronal activity on pathological protein spreading patterns is becoming increasingly clear and provokes the need for mechanistic, mathematical modeling of the bidirectional relationship between disease progression and neuronal activity. Building our model from mechanistic principles from the neuroscientific literature, we provide a simple mathematical model of this relationship. Importantly, the heterodimer-oscillator provides falsifiable hypotheses on the nature of prion-like spreading; protein spreading patterns follow a neuronal activity gradient and more extreme gradients push the brain towards a pathological state. We have also demonstrated that the heterodimer-oscillator indeed alters the tau staging patterns when simulated on a human brain connectome. It is not uncommon for patients to deviate from the stereotypical Braak staging patterns, and the heterodimer-oscillator may thus provide a mechanistic explanation for such aberrations. Future work is needed to establish the predictive power and ramifications of the heterodimer-oscillator in applications to neurodegenerative disease modeling.

## 6 Author contributions

All authors have contributed equally to the conception, theory, and writing of this work. All simulations and implementations were performed by C.G.A.

## 8 Data Accessibility

Supporting data for this research will be given upon request.

## 9 Ethics

The authors declare that they have no competing interests.

## A Coefficients of the cubic

The toxic solution for the skewed heterodimer model is given by the real positive solution (when it exists) of a cubic equation for the toxic fixed point (*v*_2_ ≠ 0) given by

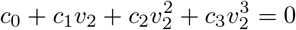

with

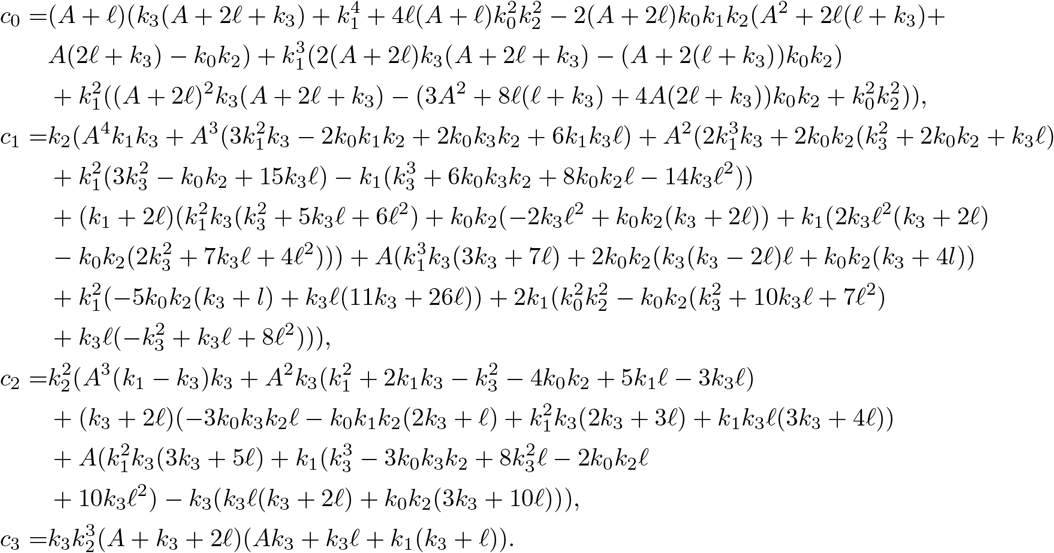

## B Critical clearance for the skewed heterodimer model

In this section, we show that the critical clearance of the 2-node skewed heterodimer model is constrained to the interval [*k*_0_*k*_2_*/k*_1_, 2*k*_0_*k*_2_*/k*_1_] and is monotonically increasing in the activity parameter *A*.

### B.1 Critical clearance bounds

In the skewed heterodimer 2-node system (Section 2.2), the healthy fixed point switches stability at a critical value for toxic clearance given by

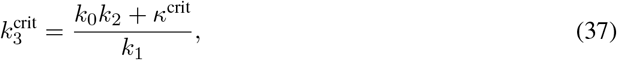

where *κ*^crit^ is given in Section 2.2. We now verify the statement 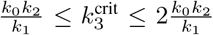 by showing that 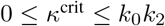. As all parameters are nonnegative the following inequality holds

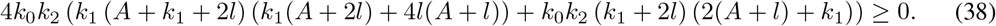

The inequality can be rewritten in terms of *s*_0_ and *s*_1_ as defined in Section 2.2, giving

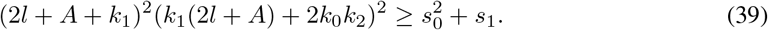

Taking the square root of both sides and rearranging gives us the desired *κ*^crit^ ≤ *k*_0_*k*_2_. As mentioned in Section 2.2, *κ*^crit^ ≥ 0 since all parameters are nonnegative. Conclusively, we have that 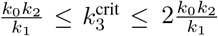.

### B.2 Monotonic dependence of critical clearance on activity

We now verify the statement that 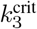 is monotonically increasing in *A* by showing that 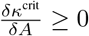. As all parameters are nonnegative, the following inequality holds

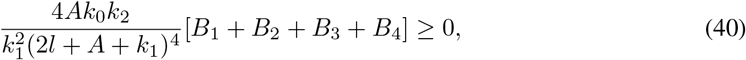

where

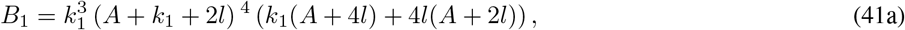

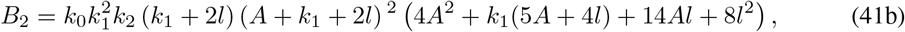

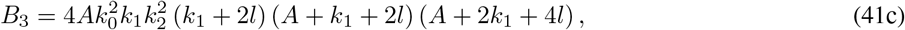

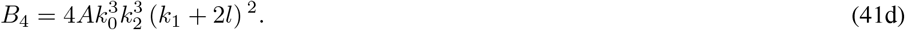

Rearranging the above inequality gives

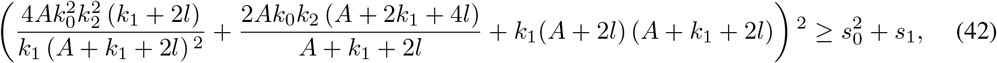

where *s*_0_ and *s*_1_ are again defined as in Section 2.2. Taking the square root of the right- and left-hand side of the above inequality, and dividing both sides by 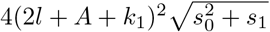 produces

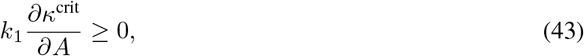

showing that 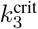 is monotonically increasing in *A*. The full expression of the partial derivative has been omitted due to its length.

